# Synthetic gene regulatory networks in the opportunistic human pathogen *Streptococcus pneumoniae*

**DOI:** 10.1101/834689

**Authors:** Robin A. Sorg, Clement Gallay, Jan-Willem Veening

**Author notes:** Correspondence to, Twitter handle: @JWVeening.

## Abstract

*Streptococcus pneumoniae* can cause disease in various human tissues and organs, including the ear, the brain, the blood and the lung, and thus in highly diverse and dynamic environments. It is challenging to study how pneumococci control virulence factor expression, because cues of natural environments and the presence of an immune system are difficult to simulate *in vitro*. Here, we apply synthetic biology methods to reverse-engineer gene expression control in *S. pneumoniae*. A selection platform is described that allows for straightforward identification of transcriptional regulatory elements out of combinatorial libraries. We present TetR- and LacI-regulated promoters that show expression ranges of four orders of magnitude. Based on these promoters, regulatory networks of higher complexity are assembled, such as logic AND and IMPLY gates. Finally, we demonstrate single-copy genome-integrated toggle switches that give rise to bimodal population distributions. The tools described here can be used to mimic complex expression patterns, such as the ones found for pneumococcal virulence factors, paving the way for *in vivo* investigations of the importance of gene expression control on the pathogenicity of *S. pneumoniae*.

## Introduction

Human pathogens and commensals reside in highly dynamic environments where they interact with host tissue, the immune system and niche competitors. *Streptococcus pneumoniae* (pneumococcus) is a prominent example of a colonizer of such complex habitats. Pneumococcus is generally found in a commensal state in the human nasopharynx; however, pneumococci can also cause disease, such as otitis media, meningitis, sepsis and pneumonia, and they are responsible for more than one million deaths per year^1,2^. To date, it remains unclear how pneumococcus switches from benign to pathogenic, and which genes are exactly involved^3,4^.

Although many studies have been dedicated to unravel key regulators and their regulons^5-11^, it remains unclear how pneumococci accurately control virulence gene expression under changing conditions. One strategy to investigate complex gene regulation relies in the approach of examining artificial regulatory networks that mimic natural networks^12^. Such approaches have yielded valuable insights on the mode of action of complex gene-regulatory networks in model organisms such as *Escherichia coli* and *Bacillus subtilis*^13-18^, but for many human pathogens these tools mostly still need to be developed.

Here, we apply synthetic biology approaches to engineer novel gene regulatory pathways in *S. pneumoniae*. A selection platform was created to enable the identification of genetic elements for specific gene expression patterns. Orthogonal transcription factors were introduced and functionalized in *S. pneumoniae*. Based on these regulators, complex gene expression networks were assembled and characterized, such as inverters, a logic AND gate and toggle switches. These tools can be applied for *in vivo* investigations of the contribution of gene expression control on the pathogenicity of *S. pneumoniae*.

## Results

### Selection and counterselection systems for *S. pneumoniae*

Synthetic gene regulatory networks are difficult to construct to date in *S. pneumoniae*, or even in model organisms, because of a lack of well-characterized individual parts, and because of limited knowledge in predicting interference between assembled parts^19-21^. To bypass these limitations, we aimed at developing an alternative strategy: the identification of regulatory elements based on selection and counterselection of genetic libraries. Parts of desired functionality, such as strong promoters, can be identified via selection, by cloning promoter libraries in front of antibiotic resistance markers ^22^. Conversely, weak promoters can be identified via counterselection^23^, and more complex transcription patterns can be found via sequential selection and counterselection.

In order to create a positive selection platform for *S. pneumoniae*, we first tested and cloned a set of five antibiotic resistance markers under the control of the Zn^2+^-inducible promoter PZ1 (Fig. 1a)^24^. As expected, cells only grew at high concentrations of antibiotics when PZ1 was induced with high concentrations of Zn^2+^. A large difference in dynamic response rate was observed between different resistance markers (Fig. 1b). Pneumococci responded to concentrations of chloramphenicol and tetracycline within one order of magnitude of Zn^2+^ induction, kanamycin spanned two orders of magnitude, and both erythromycin and trimethoprim showed a response range of more than three orders of magnitude (Fig. 1b). Since trimethoprim selection worked less well on plates and is furthermore prone to lead to emergence of spontaneously resistant cells (by SNPs in *dhfR*), we chose erythromycin and its marker *erm* as positive selector.

**Figure 1.**
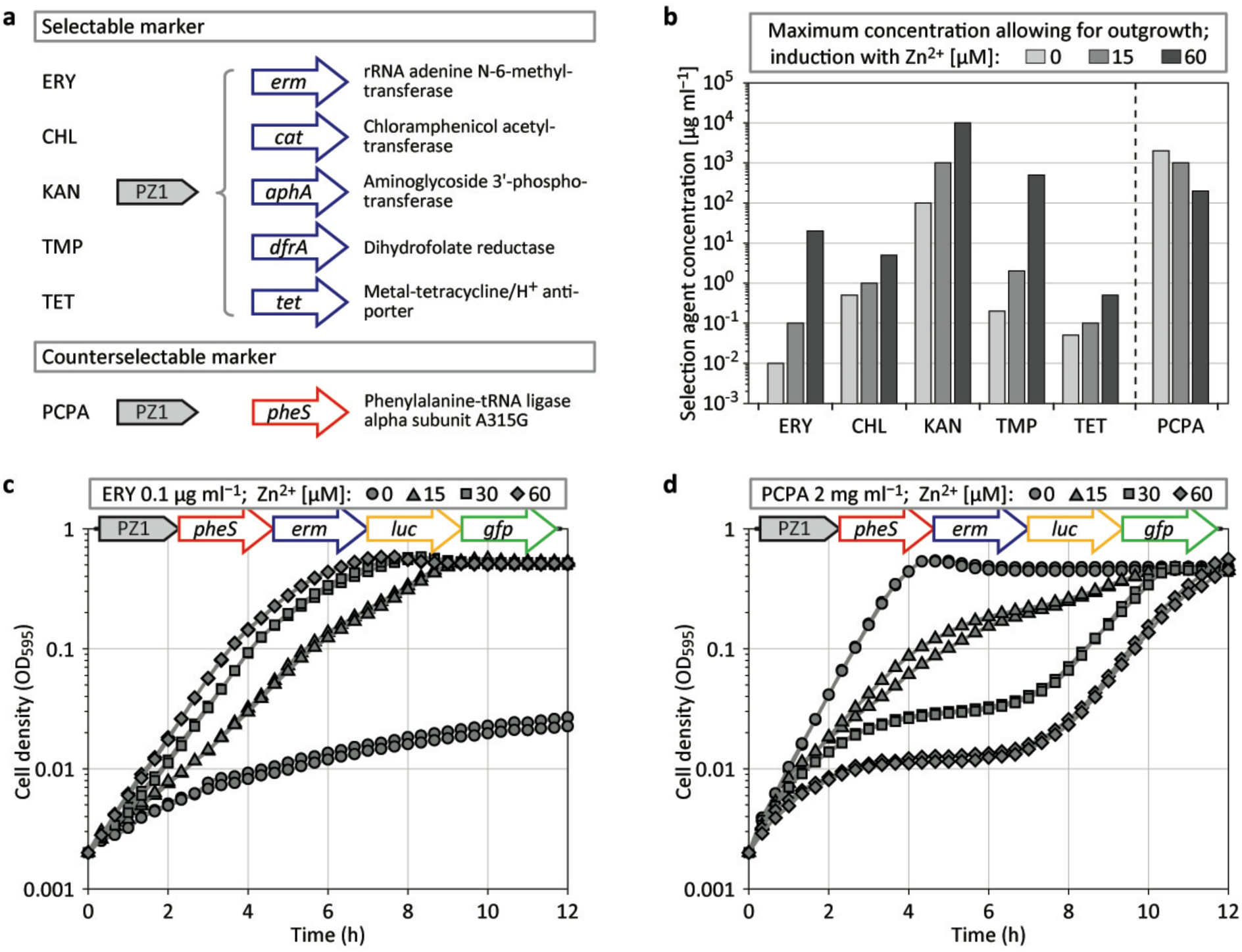
Selection marker characterization. (**a**) Schematic representation of five selectable and one counterselectable marker, including a description of their enzymatic activities, integrated into the *S. pneumoniae* D39V genome at the *amiF* (CEP) locus under the control of the Zn^2+^-inducible promoter PZ1; ERY, erythromycin; CHL, chloramphenicol; KAN, kanamycin; TMP, trimethoprim; TET, tetracycline; PCPA, *para*-chlorophenylalanine. (**b**) Determination of the maximum concentration allowing for outgrowth (starting from OD_595_ 0.002, reaching OD_595_ 0.2 or higher, within 10 h for antibiotics, and within 5 h for PCPA) of strains harboring selection and counterselection marker constructs, in dependency of Zn^2+^ induction; cultures were tested in duplicate with concentration series that doubled the applied dosage in each consecutive step (one order of magnitude was split into three steps of similar size: 1, 2, 5, 10); in the case of PCPA the upper limit of 2 mg ml^−1^ was the highest concentration tested because of solubility limitations. (**c, d**) Plate reader assay sets in duplicate measuring cell density (OD_595_) of *S. pneumoniae* D-PEP7PZ1 (see scheme; *luc*, luciferase; *gfp*, green fluorescent protein) growing at different induction levels of PZ1, in the presence of 0.1 μg ml^−1^ ERY (**c**), or in the presence of 2 mg ml^−1^ PCPA (**d**).

To date, and to our knowledge, there is only one counterselection system described for *S. pneumoniae* called Janus^25^. Janus cloning relies on a streptomycin-resistant strain that becomes susceptible when the wild type allele of the ribosomal gene *rpsL* is expressed. Janus was furthermore extended by adding the *B. subtilis* gene *sacB* (Sweet Janus)^26^ which confers sucrose sensitivity via an unknown mechanism^27^; however, counterselection with *sacB* was never shown to work independently of the Janus context in *S. pneumoniae*^26^. The disadvantage of Janus cloning is the requirement of a genetic background that carries a mutated *rpsL* allele. The *pheS* counterselection system of *E. coli* does not rely on a mutated genetic background^28^. Expression of the mutated phenylalanine-tRNA ligase PheS^A294G^ allows for the incorporation of a toxic analog of phenylalanine, called *para*-chlorophenylalanine (PCPA), into proteins which in turn causes growth retardation. We introduced this system into *S. pneumoniae* by expressing the pneumococcal equivalent of *E. coli* PheS^A294G^, *S. pneumoniae* PheS^A315G^, under control of PZ1 (Fig. 1a), and furthermore under control of a strong constitutive promoter. In the presence of PCPA the expression of PheS^A315G^ indeed led to reduced growth, both in liquid culture (Fig. 1b) and on agar plates (SFig. 1; see also Methods).

After the identification of a suitable selection system (*erm*/erythromycin) and a counterselection system (*pheS*/PCPA) we combined the two markers, together with *luc* (firefly luciferase), allowing for the analysis of gene expression at the population level, and with *gfp* (green fluorescent protein), allowing for single-cell analysis. BglBrick cloning was used to assemble genes in a consecutive manner into the vector pPEP^24^ (see Methods). The resulting plasmid, named pPEP7, was first tested by placing the polycistronic expression cassette under control of the Zn^2+^-inducible promoter PZ1. Plasmid pPEP7PZ1 (harboring PZ1-*pheS*-*erm*-*luc-gfp*) was transformed into *S. pneumoniae* D39V^29^, resulting in strain D-PEP7PZ1. As shown in Figure 1c, when strain D-PEP7PZ1 was grown in the presence of 0.1 μg ml^−1^ erythromycin, a clear Zn^2+^-dependent growth profile was observed. In contrast, when D-PEP7PZ1 was grown in presence of 2 mg ml^−1^ PCPA, the dose–response relationship of Zn^2+^ induction was inverse; the absence of Zn^2+^ allowed for uninhibited growth while high Zn^2+^ levels resulted in growth arrest (Fig. 1d). However, after a lag period of approximately 6 h, Zn^2+^-induced cells restarted to grow (Fig. 1d) suggesting the rapid emergence of mutants.

### Efficient selection of a wide range of constitutive synthetic promoters

We tested the ability of the new selection platform to identify genetic elements of desired function by screening promoter libraries. We used the strong synthetic promoter P2^24^ as a template for four distinct promoter libraries, containing randomized sequences in: (i) the UP element (UP, position −58 to −36 relative to the transcription start site^30^, resulting in 4^23^ = 7.0×10^13^ potential combinations); (ii) the core region (CORE, the 17 nucleotides between the −35 and the −10 hexamers, 4^17^ = 1.7×10^10^ potential combinations); (iii) the −35 and −10 hexamers (TATA, only specific sequence variations were allowed^31^, resulting in 2^6^ = 64 potential combinations); (iv) the proximal region (PROX, in this case the 14 nucleotides immediately downstream of the −10 hexamer, 4^14^ = 2.7×10^8^ potential combinations) (Fig. 2a).

**Figure 2.**
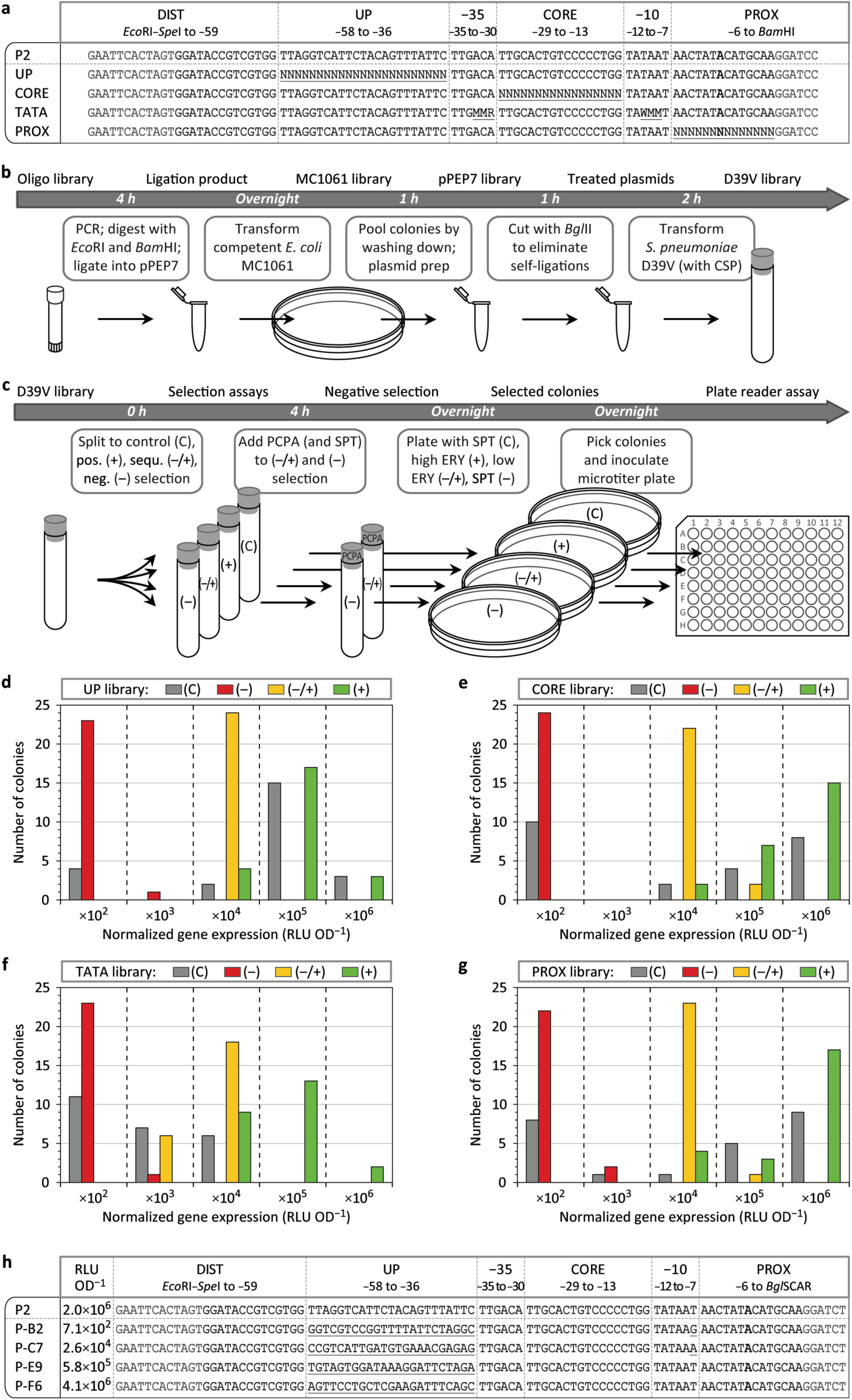
Construction and selection of promoter libraries. (**a**) Sequence of the strong constitutive promoter P2 and of four corresponding promoter libraries harboring randomized sections; DIST, distal region; UP, UP element; −35, −35 hexamer; CORE, core region; −10, −10 hexamer; PROX, proximal region; transcription start sites are shown in bold, restriction sites are shown in dark grey, randomized sequences are underlined; N=A/C/G/T, M=A/C, R=A/G, W=A/T. (**b**) Work flow of the promoter library construction, starting from oligonucleotide libraries via pPEP7 plasmid libraries to *S. pneumoniae* D39V libraries; CSP, competence-stimulating peptide. (**c**)Work flow of selection assays of D39V libraries and the identification of the promoter strengths of individual strains; C, control; (–), negative selection; (–/+), sequential selection; (+), positive selection; PCPA, *para*-chlorophenylalanine, 2 mg ml^−1^; SPT, spectinomycin, 100 μg ml^−1^; ERY, erythromycin, high: 5 μg ml^−1^, low: 0.05 μg ml^−1^. (**d**-**g**) Gene expression strength (luminescence from luciferase expression) of P2 promoter variants from 24 randomly selected colonies per selection condition of the UP library (**d**), the CORE library (**e**), the TATA library (**f**), and the PROX library (**g**); ×10^2^ etc., normalized luminescence between 1.0×10^2^ and 9.9×10^2^ RLU OD^−1^ (relative luminescence units per optical density at 595 nm) (see also Methods). (**h**) Sequence and expression strength of four promoters of the UP library (in comparison to P2), derived from negative selection (P-B2), sequential selection (P-C7), and positive selection (P-E9 and P-F6); transcription start sites are shown in bold, restriction sites are shown in dark grey, sequence deviations are underlined.

Promoter libraries were constructed by extension of two oligonucleotides that overlapped approximately 20 base pairs, with one oligonucleotide containing a randomized section (Fig. 2a, b). Double-stranded promoter constructs were integrated into pPEP7 via BglBrick assembly and sub-cloned in *E. coli* MC1061. Approximately 3500 colonies were pooled per library, and plasmid DNA was isolated, cut with *Bgl*II to eliminate self-ligations (in this case the restriction site disappeared upon promoter integration), and the resulting four pPEP7 libraries were transformed into *S. pneumoniae* D39V (Fig. 2b; see also Methods).

D39V libraries underwent four different selection treatments (Fig. 2c). Cells were plated either with spectinomycin (resistance marker of the pPEP backbone) to obtain all possible promoters (control), or with a high concentration of erythromycin to obtain strong promoters (positive selection). Alternatively, transformed cultures were first treated with PCPA for 4 h (and additional spectinomycin to inhibit non-transformed cells) and subsequently plated either with a low dosage of erythromycin for intermediate promoters (sequential selection), or with spectinomycin for weak promoters (negative selection). After overnight incubation, cells from 24 individual colonies per selection condition were analyzed by cultivation in microtiter plates. Luminescence from luciferase expression served as readout of the promoter strength.

Transformants originating from the control treatment demonstrated a wide range of promoter strengths, as measured by luminescence (Fig. 2d–g). In case of the UP, CORE and PROX libraries, at least half of the control transformants showed high levels of luminescence (1.0×10^5^ to 9.9×10^6^ RLU OD^−1^, relative luminescence units per optical density at 595 nm) (Fig. 2d, e, g). In contrast, random sampling of isolates from the TATA library did not yield promoters with similar strength compared with the template promoter, which carries the consensus TATAAT sequence, confirming the importance of the canonical −35 and −10 sequences for functional promoters (Fig. 2f). Importantly, promoters of a desired strength (strong, intermediate or weak) could be selectively enriched from all four promoter libraries (Fig. 2d–g). For example, when selecting with PCPA, only transformants showing low levels of luminescence were recovered, while selection in the presence of erythromycin led to the recovery of transformants with high gene expression activity (Fig. 2d–g). To examine whether the libraries were diverse in sequence, four promoters of the UP library (one weak, one intermediate, and two strong ones) were sequenced. Indeed, all UP element sequences deviated from the P2 template promoter (Fig. 2h). Interestingly, both the weak and the intermediate promoters contained an additional sequence deviation within the −10 hexamer, suggesting that random sequence variations within the UP element did not frequently result in an impairment of the promoter activity of P2.

### TetR- and LacI-regulated promoters

The above-mentioned results showed that our cloning vector and our selection platform could be successfully applied to identify constitutive promoters of desired strength. Next, we sought to identify controllable promoters, from which transcription can be induced by the exogenous addition of small molecules. To date, there are only few inducible systems available for *S. pneumoniae*, showing different drawbacks for specific applications. These systems are either based on pneumococcal regulators and they are thus not orthogonal^24,32-34^, or they are regulated by peptides and thus require complex, membrane associated uptake and signaling machineries^35,36^, or they show a limited dynamic range^37^. We aimed at introducing orthogonal transcription factors into *S. pneumoniae* that are regulated by small diffusible molecules, and that enable a large dynamic range. The most commonly used and best characterized bacterial regulators are the TetR and LacI repressors, which bind in the form of dimers to operator sites (*tet*O and *lac*O, respectively) consisting of 19 to 21 base pair-long DNA sequences with dyad symmetry. TetR originates from the tetracycline-resistance operon encoded in Tn10 of *E. coli* and responds to the antibiotic tetracycline^38,39^ while LacI represses the *lac*-operon and responds to the sugar allolactose^40^. These compounds interact with their corresponding repressors and trigger conformational changes that dramatically decrease the binding affinity for operator sites. Importantly, non-toxic and non-degradable inducer molecules to these repressors exist, ATc (anhydrotetracycline) in the case of TetR, and IPTG (isopropyl β-D-1-thiogalactopyranoside) in the case of LacI.

We codon-optimized *tetR* and *lacI* for expression in *S. pneumoniae* D39V and integrated them, together with an optimal pneumococcal ribosome binding site^24^, upstream of the multiple cloning site of pPEP, under control of the previously identified strong constitutive promoter PF6 (Fig. 2h). The resulting plasmids pPEP8T and pPEP8L harbor *luc* and *gfp* in the BglBrick cloning site for gene expression analysis (Fig. 3b, c). Oligonucleotide libraries used to identify constitutive promoters (Fig. 2) are less helpful for selecting TetR- and LacI-repressed promoters because the number of spacer nucleotides that separate operator sites from critical promoter sequences, such as the −10 hexamer, cannot be easily randomized using standard oligonucleotide synthesis. In this case, we took a more directed approach and placed operator sites in altering positions within the core and the proximal region of P2, which were found to be tolerant for sequence variations (Fig. 2e, g). The sequences of five promoters for TetR (PT) and five promoters for LacI (PL) are shown in Figure 3a. PT and PL promoters were cloned in front of *luc* and *gfp* into pPEP8T and pPEP8L, and plasmids were transformed into *S. pneumoniae* D39V. Transformants were grown and analyzed by microtiter plate reader assay in duplicate with one well serving as a control for full promoter repression (no inducer), and one well containing saturating concentrations of inducer molecules for maximum induction.

**Figure 3.**
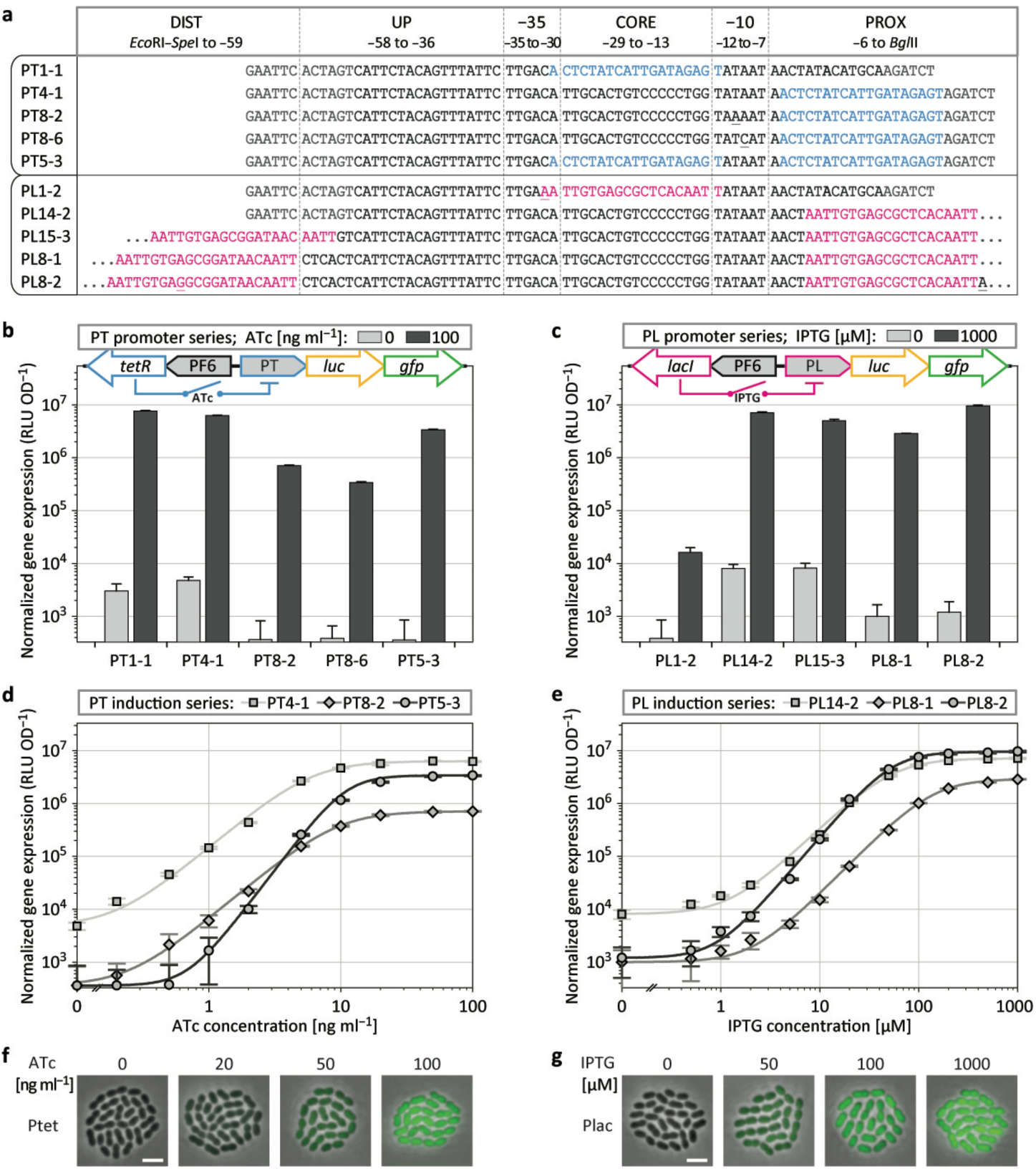
ATc- and IPTG-inducible promoters. (**a**) Sequence of TetR-repressed PT promoters and LacI-repressed PL promoters, based on the constitutive promoter P2, with TetR operator sequences in light blue and LacI operator sequences in magenta; transcription start sites are shown in bold, restriction sites are shown in dark grey (not shown for most LacI promoters because of space limitations), sequence deviations (originating from degenerate oligos or from spontaneous mutations during cloning) are underlined. (**b, c**) Luminescence from luciferase expression, driven by PT promoters (**b**) and PL promoters (**c**), without induction and with maximum induction, integrated at the *amiF* locus together with the strong constitutive promoter PF6 driving regulator expression; ATc, anhydrotetracycline; IPTG, isopropyl β-D-1-thiogalactopyranoside; error bars see Methods. (**d, e**) Induction series of selected PT promoters with ATc (**d**) and of selected PL promoters with IPTG (**e**), measured by normalized luminescence; error bars and fit curves see Methods. (**f, g**) Overlay of phase contrast and fluorescence microscopy of pneumococcal cells expressing GFP driven by PT5-3 (Ptet) in dependency of ATc induction (**f**) and driven by PL8-2 (Plac) in dependency of IPTG induction (**g**); scale bar, 2 μm.

Within the PT promoter series, a single *tet*O site (*tet*O_1_)^41^ was placed either into the core region (PT1-1) or into the proximal region (PT4-1) of P2, which gave rise to similar results, with expression values for the induced and the repressed state within approximately three orders of magnitude (Fig. 3a, b). For PT5-3, two operator sites were placed both into the core and into the proximal region, resulting in a gene expression activity of 3.4×10^6^ RLU OD^−1^ in the presence of ATc, and 3.6×10^2^ RLU OD^−1^ in the absence of ATc (indistinguishable from control cultures without luciferase), and thus giving rise to a dynamic expression range of approximately four orders of magnitude (Fig. 3b).

We furthermore analyzed −10 sequence variants of PT4-1, harboring TAAAAT in the case of PT8-2, and TATCAT in the case of PT8-6 (Fig. 3a). Both the induced and the repressed expression values of PT8-2 were downshifted one order of magnitude as compared to PT4-1, demonstrating that promoter leakiness in the repressed state can be reduced by decreasing the overall expression strength (Fig. 3b). PT8-6 showed even lower expression in the induced state as compared to PT8-2; luminescence of the repressed state, however, did not decrease any further because the lower detection limit for luciferase expression was reached (Fig. 3b). The expression curve of an induction series of PT8-2, as compared to PT4-1, was found to be similar but downshifted. Interestingly, PT5-3, harboring two *tet*O sites, showed a more hypersensitive dose-response relationship for ATc induction as compared to PT4-1 and PT8-2 that harbor only one *tet*O site (Fig. 3d).

For the PL promoter series, the positioning of *lac*O sites (*lac*O_sym_)^42^ within the core region (PL1-2) resulted in weak expression in the induced state, presumably because of the introduction of a −35 sequence variation (Fig. 3a, c). In contrast, positioning in the proximal region (PL14-2) resulted in a strong expression of 7.1×10^6^ RLU OD^−1^ when adding IPTG. Luminescence of repressed cultures was not completely suppressed and gave rise to a measurement of 8.1×10^3^ RLU OD^−1^ (Fig. 3c). LacI repressors are known to be able to tetramerize by binding two operator sites that are in proximity to one another^43^. The spacing between these two *lac*O sites is critical because repressor molecules need to face each other in order to loop DNA and form stable LacI tetramer–DNA complexes^44^. Placing an additional *lac*O site (*lac*O_1_)^45^ at the distal region (72.5 bps upstream of the first operator site, as measured from center to center of the *lac*O sites) did not lead to an improved repression, and luminescence levels were similar as compared to PL14-2 that harbors only one *lac*O site. However, when shifting the second operator four base pairs further upstream, we could observe an 8–fold reduction of luminescence in the repressed state as compared to PL14-2; unfortunately, this also led to a 2–fold reduction in the induced state (Fig. 3c). Serendipitously, another clone with a double *lac*O site was isolated, called PL8-2, which differs from PL8-1 by two spontaneous insertions, one additional nucleotide in the distal operator site and one additional nucleotide in the proximal region (Fig. 3a). PL8-2 showed the largest induction range of the PL promoter series, spanning four orders of magnitude, with an expression strength of 9.5×10^6^ RLU OD^−1^ in the induced state and 1.2×10^3^ RLU OD^−1^ in the repressed state (Fig. 3c). PL8-2 also showed the strongest hypersensitive response towards IPTG induction (Fig. 3e).

To examine controllable gene expression at the single-cell level, strains D-PEP8T5-3 and D-PEP8L8-2, harboring promoters with the highest dynamic range (promoters PT5-3, hereafter called Ptet, and PL8-2, hereafter called Plac) were grown at varying concentrations of ATc and IPTG, and GFP expression was visualized by fluorescence microscopy. A clear dose– response relationship was found within the induction series of each strain (Fig. 3f, g). Together, these results suggest that Ptet and Plac display a dynamic induction range of four orders of magnitude, making them the best controllable promoters currently available for *S. pneumoniae*. Indeed, our laboratory has successfully used Plac in several studies to accurately express transcription of various genes in *S. pneumoniae*^46,47^. Recently, another group also established a LacI-based IPTG-inducible system for use in *S. pneumoniae*^48^, further demonstrating the demand for such small-molecule orthogonal inducible systems in this organism.

### Inducible systems performing logical operations

To build gene regulatory networks of higher complexity, individual parts need to work in a robust manner, independent of their genomic location. To test whether the here-described repressor systems can be expressed from an ectopic locus, and thus distal to the promoters which they regulate, we integrated the TetR and LacI repressors at the non-essential *prsA* locus^49^ (see Methods) (Fig. 4b). Genes of interest under the control of Ptet or Plac were integrated at the *amiF* (CEP) locus via the pPEP9 series of plasmids, which differs from pPEP8T and pPEP8L by omitting the repressors. Gene expression activity of Ptet (strain D-T-PEP9Ptet) and Plac (strain D-L-PEP9Plac) was similar as compared to strains harboring the repressor cassette at the *amiF* locus (strains D-PEP8T5-3 and D-PEP8L8-2, respectively) (Figs 3d, e, and 4c, d).

**Figure 4.**
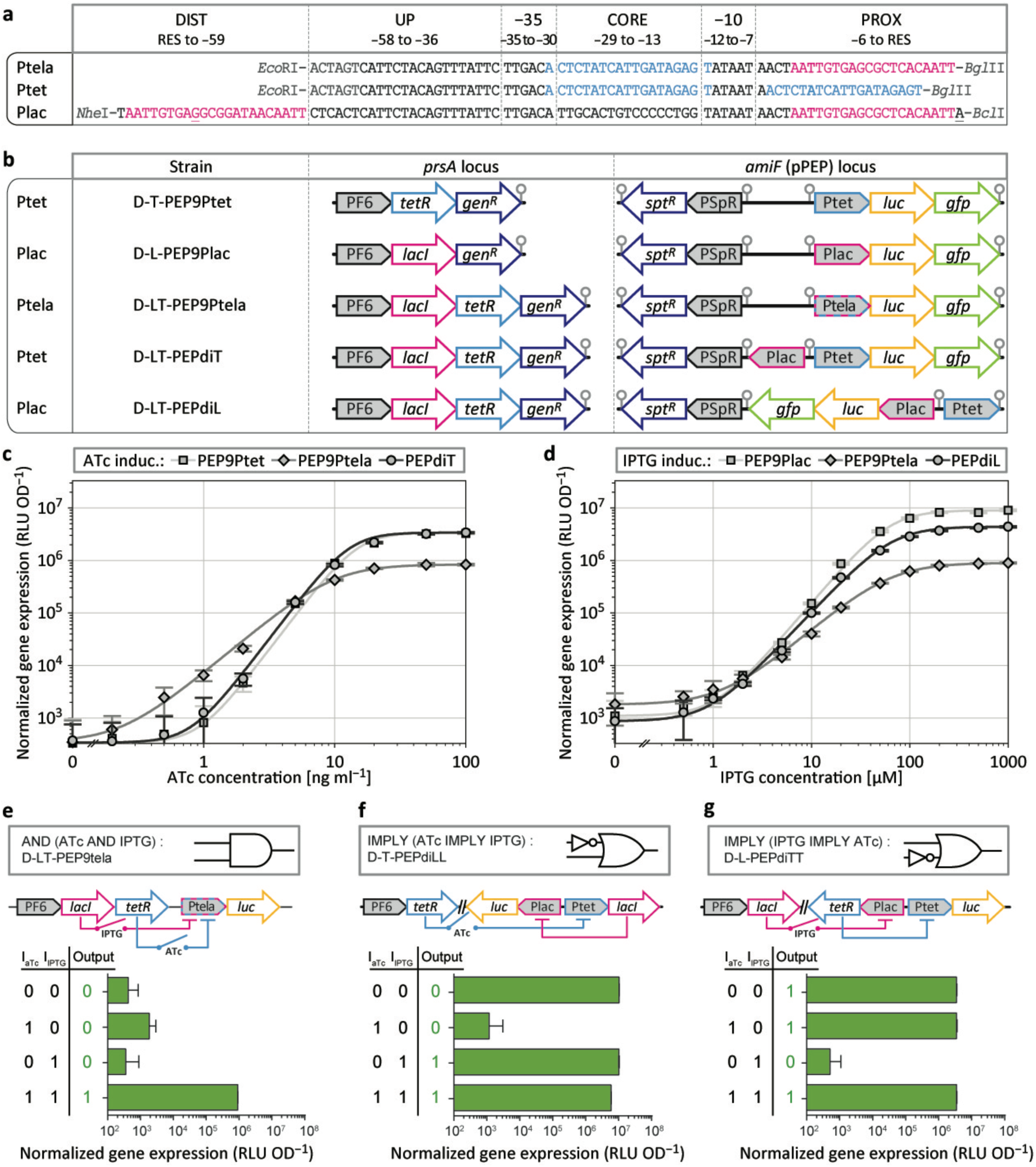
Construction of inverters, amplifiers and a logic AND gate. (**a**) Sequence and restriction sites of the TetR+LacI double-repressed promoter Ptela, and Ptet (renamed PT5-3) and Plac (renamed PL8-2) in the context of the double-inducible system PEPdi, with TetR operator sequences in light blue and LacI operator sequences in magenta; transcription start sites are shown in bold, restriction sites are indicated in dark grey. (**b**) Schematic representation of gene expression regulation constructs, with the regulators expressed from the *prsA* locus and the genes of interest expressed from the *amiF* locus; *gen*^*R*^, gentamicin resistance marker; *spt*^*R*^, spectinomycin resistance marker; grey circles indicate transcription terminators. (**c, d**) Induction series of TetR-regulated promoters with ATc (**c**) and of LacI-regulated promoters with IPTG (**d**) of the strains shown in **b**, measured by luminescence; for Ptela induction series with ATc, 1000 μM IPTG was added to de-repress LacI; for Ptela induction series with IPTG, 100 ng ml^−1^ ATc was added to de-repress TetR; error bars and fit curves see Methods. (**e, f, g**) Boolean logical operators, expressing luciferase in the absence of inducer and during induction with ATc (100 ng ml^−1^), IPTG (1000 μM) and ATc + IPTG, from Ptela (**e**), Plac (**f**), and Ptet (**g**).

More complex programming of gene expression, as for example Boolean logic gates, requires the integration of multiple signals into one single response^50,51^. We wondered whether a P2-based promoter could be used to create a logic AND gate for external induction, requiring both ATc and IPTG. To test this, the synthetic promoter Ptela was constructed that contains a *tet*O site within the core region and a *lac*O site within the proximal region (Fig. 4a). Strain D-LT-PEP9Ptela, which drives *luc* from Ptela and expresses both LacI and TetR from the *prsA* locus, was found to require both ATc and IPTG to highly express luciferase (Fig. 4c, d, e). LacI repression on its own (with ATc but without IPTG) was not enough to completely shut down Ptela activity (Fig. 4e). However, the absence of ATc, and thus TetR repression alone, was enough to decrease luminescence below the detection limit even in the presence of 1 mM of IPTG (Fig. 4e).

The results above show that both TetR and LacI can be functional within the same cell. Next, we wondered whether the ATc- and IPTG-inducible systems could be used in parallel and independent from one another. We therefore generated the double-inducible integration plasmid pPEPdi, with promoter Ptet positioned in the BglBrick cloning site, and promoter Plac in an upstream inverse position within the terminator-insulated BglBrick transfer site^24^. Note that Plac, in the context of pPEPdi, harbors different flanking restriction sites, with *Nhe*I upstream and *Bcl*I downstream of the promoter sequence (Fig. 4a). Two variants of pPEPdi were tested in strains expressing both LacI and TetR at the *prsA* locus: D-LT-PEPdiT driving *luc* and *gfp* from Ptet, and D-LT-PEPdiL driving *luc* and *gfp* from Plac (Fig. 4b). Ptet induction in this context was found to closely match the values obtained with PT5-3, without any observable interference from the additionally present LacI (Fig. 4c). Plac expression, in contrast, deviated from the corresponding PL8-2 pattern, with a 2–fold decreased maximum luminescence in the presence of high concentrations of IPTG (Fig. 4d). Weaker luminescence signals from Plac in the double-inducible system, as compared to PL8-2, could originate from a decreased promoter activity caused by the inverse reading orientation (into the direction of DNA replication) or from the sequence deviation in the proximal region (*Bcl*I instead of the *Bgl*II site). Alternatively, the translation efficiency might be decreased because of the alteration within the 5’ UTR. Nevertheless, the double-inducible system showed to work without interference between the two regulators.

With the TetR- and LacI-regulated systems in place, we wondered if we could construct a system where the expression of one repressor is controlled by the activity of the other repressor, giving rise to IMPLY gates (Fig. 4f, g). To do so, we controlled the amount of LacI via Ptet induction (strain D-T-PEPdiLL; scheme in Fig. 4f), and the amount of TetR via Plac induction (strain D-L-PEPdiTT; scheme in Fig. 4g). Indeed, both strains D-T-PEPdiLL and D-L-PEPdiTT were found to act as effective IMPLY gates. Interestingly, while Ptet showed the same maximum expression in the absence of TetR as in the presence of fully de-repressed TetR, in the case of Plac, luminescence was 2–fold higher in the absence of LacI as compared to fully de-repressed LacI. This observation confirms the findings of a previous study with *E. coli* showing that de-repressed LacI does not completely lose its affinity for *lac*O sites^52^.

### Construction and characterization of single-copy toggle switches

Heterogeneous expression of virulence factors within bacterial populations, such as the polysaccharide capsule in *S. pneumoniae*, may contribute importantly to pathogenicity^53-55^. To be able to improve our understanding of such expression patterns, we aimed at engineering synthetic regulatory networks that are bistable, and that can thus give rise to bimodal population distributions^12^. A classic example of such a network is the so-called toggle switch, in which two transcription regulators repress the expression of each other^14^. The *prsA* site was used to integrate a genetic toggle switch into the pneumococcal genome (Fig. 5a). Note that that there are only few descriptions of a single-copy chromosomally integrated synthetic toggle switch, in contrast to toggle switches on replicating plasmids^56^.

**Figure 5.**
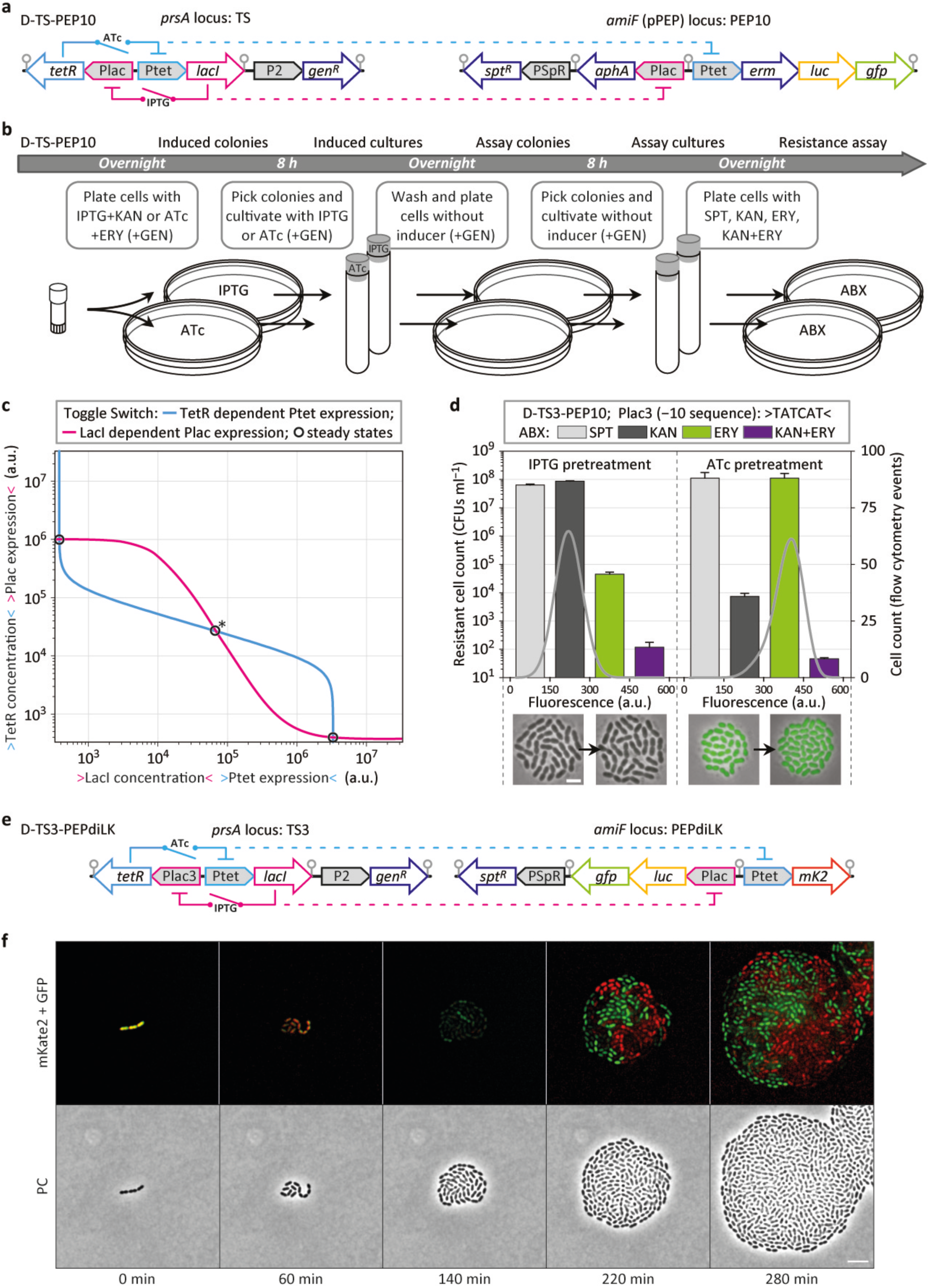
Identification of toggle switches. (**a**) Schematic representation of strain D-TS-PEP10 containing a transcriptional toggle switch at the *prsA* locus (TS) and genes of interest at the *amiF* locus (PEP10); *gen*^*R*^, gentamicin resistance marker; *spt*^*R*^, spectinomycin resistance marker; *aphA*, kanamycin resistance marker; *erm*, erythromycin resistance marker; grey circles indicate transcription terminators. (**b**) Work flow of D-TS-PEP10 induction and the subsequent identification of switching events; IPTG, 1000 μM; ATc, 100 ng ml^−1^; GEN, gentamicin, 20 μg ml^−1^; SPT, spectinomycin, 100 μg ml^−1^; KAN, kanamycin, 500 μg ml^−1^; ERY, erythromycin, 1 μg ml^−1^; KAN+ERY, kanamycin 500 μg ml^−1^ and erythromycin 1 μg ml^−1^; ABX, antibiotics. (**c**) Overlay of the fit curves corresponding to TetR-dependent Ptet expression (light blue) and LacI-dependent Plac3 expression (magenta) to indicate stable states (circles) and the threshold (circle with asterisk) of the toggle switch. (**d**) Number of resistant cells that were able to form colonies (CFUs ml^−1^, colony forming units per 1 ml cell culture at OD_600_ 0.1) from cultures derived after plating without inducer (average and s.e.m. of experimental duplicates are shown), and flow cytometry analysis of these cultures measuring the fluorescence intensity of 10^4^ cells (grey lines, displaying output levels #0 to #600 of a 10-bit channel, arbitrary units, see also Methods); underneath, an overlay of phase contrast and fluorescence microscopy of D-TS3-PEP10 cells are shown, with cells originating from induced cultures on the left and cells originating from cultures after plating without inducer shown on the right side of the arrow; scale bar, 2 μm; D-TS3-PEP10, Plac3 (−10 sequence): TATCAT. (**e**) Schematic representation of strain D-TS3-PEPdiLK, harboring toggle switch 3 (TS3) at the *prsA* locus and reporter genes at the *amiF* locus; *mK2*, mKate2; grey circles indicate transcription terminators. (**f**) Still images (phase contrast and fluorescence microscopy) of a time-lapse experiment of D-TS3-PEPdiLK cells previously treated with 100 ng ml^−1^ ATc + 50 μM IPTG growing on a semi-solid surface without inducer; scale bar, 5 μm.

Toggle switch 1 (TS1) harbors the promoters Ptet (identical to PT5-3; with restriction sites *Xba*I upstream and *Ase*I downstream) driving *lacI*, and Plac (identical to PL8-2; with restriction sites *Xba*I upstream and *Bgl*II downstream) driving *tetR* (Fig. 5a). Reporter genes were integrated into the *S. pneumoniae* D39V genome at the *amiF* locus. Based on pPEPdi, pPEP10 was created, with the erythromycin resistance marker *erm, luc* and *gfp* driven from Ptet, and the kanamycin resistance marker *aphA* driven from Plac (Fig. 5a). Separating the toggle switch from reporter genes increased the robustness of the system, and it furthermore allowed for straightforward replacement of reporter genes. Toggle switch strains were triggered, either with IPTG or with ATc, by plating and overnight incubation, followed by 8 h cultivation in liquid medium in the presence of inducer (to allow for the establishment of stable expression equilibria; Fig. 5b). Next, induced cultures were re-plated and re-grown for 8 h in liquid medium without inducer to allow for the settlement of gene expression at stable states, and for switching events to occur (Fig. 5b).

When assuming that Ptet and Plac expression in the toggle switch context result in identical patterns as found for the double-inducible system, then the TetR regulated expression of Ptet and the LacI regulated expression of Plac can be combined into one plot, displaying the gene expression regulation of the toggle switch (SFig. 2). Intersections of the two expression curves represent steady states in which gene expression activity of the two promoters and the concentration of the two repressors are in equilibrium. Three steady states emerge, with the outer two representing stable states, and the middle one representing the threshold of the bistable switch. The proximity of this threshold to one of the stable states indicates that naturally occurring fluctuations within cells might be sufficient to trigger switching events. Fluctuations can originate, amongst others, from unequal distributions of repressor molecules after cell division or from prolonged promoter accessibility after spontaneous repressor dissociation.

In the case of the promoter pair Ptet and Plac (TS1), the predicted positioning of the steady states indicates that cells containing high levels of TetR and low levels of LacI are more likely to remain in their current state than cells containing low levels of TetR and high levels of LacI (SFig. 2). We therefore created two additional toggle switches, TS2 and TS3, by modifying the −10 sequence of Plac (TS2, Plac2 −10 sequence TAAAAT; TS3, Plac3 −10 sequence TATCAT) with the goal of finding bistability patterns with similar spacing between the threshold and the two stable states. Deviations of the canonical −10 sequence were previously shown to downshift the expression curve of an induction series (10–fold for TAAAAT, and presumably 20–fold for TATCAT; see Fig. 3b, d).

The three resulting strains D-TS1-PEP10, D-TS2-PEP10 and D-TS3-PEP10 (Fig. 5b and SFig. 2) were analyzed for resistance towards: (i) spectinomycin (resistance marker of PEP), indicating the total number of viable cells; (ii) kanamycin, indicating the number of cells with low levels of LacI (and presumably high levels of TetR; from here on T-state); (iii) erythromycin, indicating the number of cells with low levels of TetR (and presumably high levels of LacI; from here on L-state); (iv) kanamycin and erythromycin, serving as a control (Fig. 5d and SFig. 3a, b). Cells of colonies that emerged in the presence of both kanamycin and erythromycin likely contained mutations. Cell populations were furthermore analyzed by flow cytometry, whereat high GFP-fluorescence indicated cells in the L-state, and low fluorescence indicated cells in the T-state (Fig. 5d and SFig. 2). Additionally, cells of both induced cultures and of cultures after re-plating without inducer were analyzed by fluorescence microscopy (Fig. 5d and SFig. 2).

Cells harboring TS1 that were previously treated with IPTG were all found in the corresponding T-state (SFig. 2). In contrast, only approximately 50 % of the TS1 cells from ATc pretreatment were found in the L-state, and the remaining half had switched to the T-state (SFig. 2). These findings matched the prediction made based on the plot in SFig. 2. In the case of TS2, cells from IPTG-pretreated cultures were also all found in the T-state (SFig. 2). For ATc pretreatment, the number of TS2 cells that had switched from the L-state to the T-state was found to be ∼ 0.5 %, and thus two orders of magnitude lower compared with TS1 (SFig. 2). Transitions from the L-state to the T-state were found to occur even less frequently in TS3 cultures, with only ∼ 0.01 % of cells showing kanamycin resistance (Fig. 5d). However, in TS3 strains, also cells of cultures that were pretreated with IPTG were found to be able to switch, in this case from the T-state to the L-state, with a frequency of ∼ 0.1 % of cells within the observed time period (Fig. 5d).

Remarkably, an additional prediction that was made based on the plot in Fig. 5c could be confirmed. When decreasing the promoter strength of Plac within the toggle switch, one would expect a lower TetR concentration in the L-state, which in turn should result in cells expressing higher levels of GFP. Indeed, flow cytometry measurements showed that the peak of fluorescence intensity shifted in the direction of the *x*-coordinate, from TS1 over TS2 to TS3, with the mode of TS1 cells found at output level #362, for TS2 cells at #388, and for TS3 cells at #398 (arbitrary units, see also Methods) (Fig. 5d and SFig. 2).

After the characterization of TS3, we attempted to identify intermediate induction levels that would allow TS3 populations to bifurcate after inducer removal. Two strains harboring TS3 were tested in parallel, one expressing luciferase from Plac (D-TS3-PEPdiLK) and the other one expressing luciferase from Ptet (D-TS3-PEPdiT) (SFig. 2). Growth cultures were adapted to different inducer concentration ranges, followed by inducer removal and cultivation without inducer (SFig. 3). Two out of six tested conditions resulted in strong luminescence of both Plac and Ptet after inducer removal, indicating that the population indeed bifurcated, with significant fractions of cells developing towards both stable states (SFig. 3).

To examine the dynamics of a population bifurcation, we induced cells of strain D-TS3-diLK with different concentrations of IPTG and ATc and analyzed single cells by fluorescence microscopy (SFig. 3). Induced cells of starting cultures (time point 0 h) showed both GFP and mKate2 fluorescence, in dependency of the initial induction level. Within 2 h after inducer removal, the fluorescence of individual cells could be detected either in the GFP- or in the mKate2-fluorescence channel, but none of the analyzed cells resulted in signals above background levels in both channels (SFig. 3). After 4 h, the stable states were consolidated. In agreement with the toggle switch architecture, at higher initial IPTG concentrations, a larger fraction of cells developed the L-state (only expressing mKate2) and fewer cells developed the T-state (only expressing GFP). Finally, we followed population bifurcations of TS3 by time-lapse microscopy (Fig. 5f). Starting from single D-TS3-diLK cells, microcolonies of phenotypically mixed progeny emerged, with individual cells fluorescing either green or red (Fig. 5f, Movie S1).

## Discussion

Here, we created and characterized a selection tool for gene expression regulation in *S. pneumoniae* by combining the counterselection marker *pheS*, the selection marker *erm*, the population reporter *luc* and the single-cell reporter *gfp*. This selection platform was used to generate and analyze libraries of constitutive pneumococcal promoters. Operator sequences for the *E. coli*-derived repressors TetR and LacI were integrated into core and proximal promoter regions that showed tolerance for sequence variations, and a set of inducible promoters was created and characterized. Based on the two promoters with the largest dynamic range, Ptet and Plac, spanning approximately four orders of magnitude, regulatory networks of higher complexity were assembled and analyzed including the successful construction of logic gates (Fig. 4) and toggle switches (Fig. 5). This emphasizes the modularity of prokaryotic promoters and provides a framework for the construction of semisynthetic promoters in *S. pneumoniae*, which respond to their original regulators, and additionally to orthogonal regulators, such as TetR or LacI.

*S. pneumoniae* represents an interesting candidate for synthetic biology applications. The straightforward uptake of exogenous DNA and stable integration into the genome via natural competence allows for stable copy numbers and tightly controlled gene expression of artificial regulatory networks. Furthermore, pneumococci have a small genome of approximately 2 million bps, and thus reduced genetic redundancy. More than 25.000 pneumococcal genome sequences are available, ranging from strains used for fundamental research, isolates from healthy carriers and clinical isolates from patients with invasive diseases. The field of clinical and fundamental research of pneumococci could in turn also profit from synthetic biology research with the organism, by receiving new tools that might help answering research questions that are difficult to address with conventional methods. It is therefore of importance to adapt tools developed for model organisms, as a proof of principle, to bacteria of interest, such as human pathogens.

To this effect, the approaches applied here allowed us to build complex gene regulatory networks, such as genetic toggle switches, into the genome of pneumococcus to accurately control gene expression. These networks could, in a next step, be used to drive virulence factors, and thus allow for *in vivo* investigations of the contribution of gene expression regulation on the pathogenicity of the pneumococcus. In addition, the approaches used here may serve as an example for synthetic biology projects in unrelated organisms.

## Methods

### Strains and growth conditions

*S. pneumoniae* D39V was used throughout^10^, and *E. coli* MC1061 was used for sub-cloning. *E. coli* competent cells were obtained by CaCl_2_ treatment^57^; transformations were carried out via heat-shock at 42°C. *S. pneumoniae* transformations were carried out with cultures at OD (600 nm) 0.1 in the presence of 1 ng ml^−1^ CSP (competence-stimulating peptide)^58^. Promoters and genes of interest were assembled in pPEP^24^ via BglBrick cloning^59^ followed by integration into the D39V genome at the *amiF* locus. Integration constructs inside the *prsA* locus^49^ (replacing base pairs 29751 to 30077) were assembled via Gibson assembly^60^ and directly transformed to *S. pneumoniae*^61^. Pneumococcal cells were cultivated in C+Y medium^62^ (pH 6.8) supplemented with 0.5 mg ml^−1^ D-luciferin for luminescence measurements, at a temperature of 37°C. Pre-cultures for all experiments were obtained by a standardized protocol, in which previously exponentially growing cells from −80°C stocks were diluted to OD 0.005 and grown until OD 0.1 in a volume of 2 ml medium in tubes that allow for direct (in tube) OD measurement. For selection assays, and to determine the number of colony-forming units, cells were plated inside Columbia agar supplemented with 3% (v v^−1^) sheep blood and incubated overnight at 37°C. For counterselection assays with *para*-chlorophenylalanine (PCPA) equal volumes of two freshly prepared stock solutions, 20 mg ml^−1^ in 1 M NaOH and 20 mg ml^−1^ in 1 M HCl, were mixed in C+Y medium or Columbia agar; control cultures with the highest resulting NaCl concentrations (without PCPA) were found to not impact pneumococcal growth behavior. Key plasmids will be made available via addgene.

### Microtiter plate reader assays

Costar 96-well plates (white, clear bottom) with a total assay volume of 300 μl per well were inoculated to the designated starting OD value. Microtiter plate reader experiments were performed using a TECAN infinite pro 200 (Tecan Group) by measuring every 10 min with the following protocol: 5 s shaking, OD (595 nm) measurement with 25 flashes, luminescence (RLU, relative luminescence units, a.u.) measurement with an integration time of 1 s. Average and s.e.m. of normalized luminescence (RLU OD^−1^) were determined between OD 0.01 and OD 0.02, based on three measurements of duplicates (and thus six data points). Fit curves for induction series were based on the 4 parameter Hill equation:

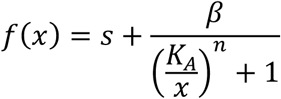

with *s*, minimum expression; *β*, maximum expression; *n*, Hill coefficient; *K*_*A*_, dissociation constant. The following parameters were found (*s*; *β*; *n*; *K*_*A*_): Fig. 3d: D-PEP8PT4-1 (4000; 6.3×10^6^; 2.1; 6.0), D-PEP8PT8-2 (360; 7.1×10^5^; 2.1; 9.3), D-PEP8PT5-3 (360; 3.4×10^6^; 3.3; 11); Fig. 3e: D-PEP8PL14-2 (8100; 7.1×10^6^; 1.8; 54), D-PEP8PL8-1 (1000; 2.9×10^6^; 1.9; 140), D-PEP8PL8-2 (1200; 9.5×10^6^; 2.2; 54); Fig. 4dc D-L-PEP9Plac (1100; 9.0×10^6^; 2.2; 63), D-LT-PEP9PLtela (1800; 9.0×10^5^; 1.6; 62), D-LT-PEPdiL (870; 4.4×10^6^; 2.0; 65); Fig. 4d: D-T-PEP9Ptet (340; 3.4×10^6^; 3.3; 15), D-LT-PEP9Ptela (350; 8.4×10^5^; 2.1; 9.7), D-LT-PEPdiT (370; 3.4×10^6^; 3.3; 13); SFig. 2a: D-L-PEPdiTT (380; 3.3×10^6^; −2.9; 7000); SFig. 2b: D-T-PEPdiLL (870; 1.0×10^7^; −1.9; 22000).

### Microscopy

A Nikon Ti-E microscope equipped with a CoolsnapHQ2 camera and an Intensilight light source were used. Microscopy was carried out by spotting cells on a 10% polyacrylamide slide containing PBS, inside a Gene Frame (Thermo Fisher Scientific) that was sealed with the cover glass to guaranty stable conditions. Images of fluorescing cells were taken with the following protocol and filter settings: 300 ms exposure for phase contrast, 1 s exposure for fluorescence at 440−490 nm excitation via a dichroic mirror of 495 nm and an emission filter at 500−550 nm. For the induction series shown in Figure 3e, f, D-PEP8T5-3 and D-PEP8L8-2 cells were pre-cultivated according to standard protocol (to OD 0.1), followed by a 100–fold dilution and regrowth to OD 0.1 in the presence of ATc and IPTG, respectively.

For time-lapse microscopy, strain D-TS3-PEPdiLK was diluted into 2 ml CY medium until a starting OD (600 nm) 0.005 and grown at 37°C until OD (600 nm) 0.3. Cells were then re-diluted into fresh CY medium supplemented with 200 μM IPTG and 100 ng μl ATc until OD (600 nm) 0.005 and grown for 8 h while keeping the OD (600 nm) < 0.05. After 8 h, 2 ml of cells at OD (600 nm) 0.05 were harvested by centrifugation 7 min at 3000xg and washed twice with 1 ml of fresh pre-warmed CY medium. Cell pellet was re-suspended into 1 ml of fresh CY medium and spotted onto a 1.2% agarose-CY pad placed into a Gene Frame and closed with a cover glass^63^. The slide was mounted onto a Leica DMi8 microscope equipped with a sCMOS DFC9000 (Leica) camera with a 100x/1.40 oil-immersion objective and an environmental chamber at 30°C, allowing cell growth. After ∼10 min, and every 20 min for 5 h, phase contrast signal was imaged using transmission light and 100 ms exposure time, and both fluorescent signals were acquired using 17% of excitation light and 400 ms exposure time with a SOLA Light Engine (lumencor). GFP signal was obtained using a GFP filter cube with 470/40 nm excitation filter (Chroma #ET470/40x) and 520/40 nm emission filter (Chroma #ET520/40m). mKate2 signal was obtained using a mCherry filter cube with excitation and emission filters at 560/40 nm and 590 nm LongPass respectively (Chroma #49017). Images were processed using LAS X (Leica) and signal was deconvolved with Huygens (SVI) software using only one iteration. Time-lapse movie was edited with Fiji software^64^.

### Flow cytometry

The fluorescence intensity of 10^4^ cells was measured by a BD FACS Canto Flow Cytometer (BD Bioscience) at medium flow, with the detectors for forward scatter and side scatter set to 200 V and 500 V, respectively. A gate was defined based on forward and side scatter measurements to exclude the recording of particles that deviated from normal *S. pneumoniae* cells. The detector for fluorescence was set to 750 V. Results shown in Fig 5d and SFig. 2 represent smoothed data (running average) of output levels #0 to #600 of a 10–bit channel (total number of 1024 output levels), whereat measurements from output levels #0 to #300 were fitted to a normal distribution for visual clarity; the detection of weakly fluorescing cells in our flow cytometer suffered from machine biases, showing periodically repeating stretches of empty reads and culmination at output level #0.

## Acknowledgements

We thank Melinde Wijers (University of Groningen, undergraduate program) for excellent technical assistance during the initial phase of this project and thank Yolanda Schaerli for useful comments on the manuscript. Work in the Veening lab is supported by the Swiss National Science Foundation (SNSF) (project grant 31003A_172861), a JPIAMR grant (40AR40_185533) from SNSF and ERC consolidator grant 771534-PneumoCaTChER.

