## Supplementary figures for "Synthetic gene regulatory networks in the opportunistic human pathogen *Streptococcus pneumoniae*"

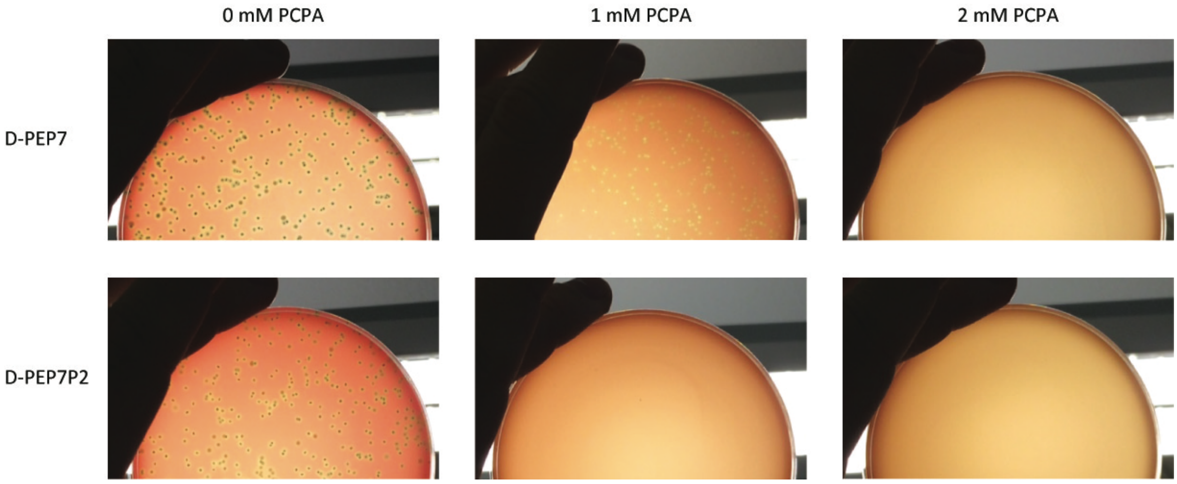


**Supplementary Figure 1 | PCPA counterselection in agar plates.** Colony formation of cells harboring *pheS* A315G, without promoter (D-PEP7) and with the strong constitutive promoter P2 (D-PEP7P2), after overnight incubation in the presence of 0, 1 and 2 mM PCPA.

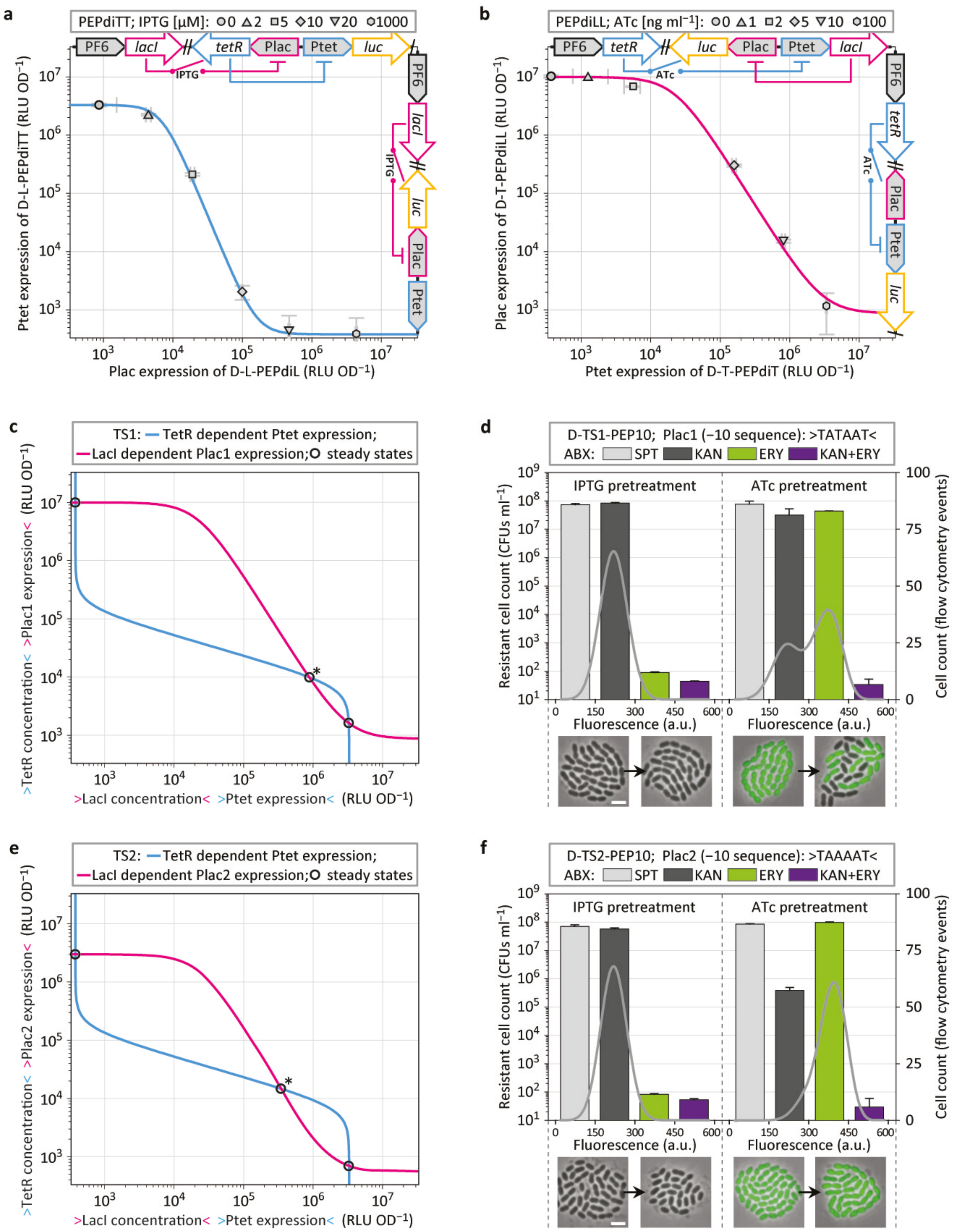

a
PEPdiTT; IPTG [μM]: 0 2 5 10 20 1000
PF6
lacI
tetR
Plac
Ptet
luc
IPTG
Ptet expression of D-L-PEPdiTT (RLU OD−1)
Plac expression of D-L-PEPdiL (RLU OD−1)
b
PEPdiLL; ATc [ng ml−1]: 0 1 2 5 10 100
ATc
Plac expression of D-T-PEPdiLL (RLU OD−1)
Ptet expression of D-T-PEPdiT (RLU OD−1)
c
TS1: TetR dependent Ptet expression;
LacI dependent Plac1 expression; steady states
>TetR concentration< >Plac1 expression< (RLU OD−1)
>LacI concentration< >Ptet expression< (RLU OD−1)
d
D-TS1-PEP10; Plac1 (−10 sequence): >TATAAT<
ABX: SPT KAN ERY KAN+ERY
IPTG pretreatment
ATc pretreatment
Resistant cell count (CFUs ml−1)
Cell count (flow cytometry events)
Fluorescence (a.u.)
e
TS2: TetR dependent Ptet expression;
LacI dependent Plac2 expression; steady states
>TetR concentration< >Plac2 expression< (RLU OD−1)
>LacI concentration< >Ptet expression< (RLU OD−1)
f
D-TS2-PEP10; Plac2 (−10 sequence): >TAAAAT<
ABX: SPT KAN ERY KAN+ERY
IPTG pretreatment
ATc pretreatment
Resistant cell count (CFUs ml−1)
Cell count (flow cytometry events)
Fluorescence (a.u.)

**Supplementary Figure 2 | Identification of toggle switches.** (**a**) IPTG induction series of D-L-PEPdiTT (upper scheme, data in the *y*-coordinate) and D-L-PEPdiL (scheme to the right, data in the *x*-coordinate), whereat luminescence of D-L-PEPdiL is proportional to the TetR concentration of D-L-PEPdiTT; error bars and fit curve see Methods. (**b**) ATc induction series of D-T-PEPdiLL (upper scheme, data in the *y*-coordinate) and D-T-PEPdiT (scheme to the right, data in the *x*-coordinate), whereat luminescence of D-T-PEPdiT is proportional to the LacI concentration of D-T-PEPdiLL; error bars and fit curve see Methods. (**c, e**) Overlay of the fit curves corresponding to TetR-dependent Ptet expression (light blue) and LacI-dependent Plac expression (magenta) to indicate stable states (circles) and the threshold (circle with asterisk) of toggle switches, with TS1 harboring Plac1 (**c**, -10 sequence: TATAAT) and TS2 harboring Plac2 (**e**, -10 sequence: TAAAAT). (**d, f**) Number of resistant cells that were able to form colonies (CFUs $ml^{-1}$, colony forming units per 1 ml cell culture at $OD_{600}$ 0.1) from D-TS1-PEP10 (**d**) and D-TS2-PEP10 (**f**) cultures derived after re-plating without inducer (average and s.e.m. of experimental duplicates are shown), and flow cytometry analysis of these cultures measuring the fluorescence intensity of $10^4$ cells (grey lines, displaying output levels #0 to #600 of a 10-bit channel, arbitrary units, see also Methods); underneath, an overlay of phase contrast and fluorescence microscopy of D-TS-PEP10 cells are shown, with cells originating from induced cultures on the left and cells originating from cultures after re-plating without inducer shown on the right side of the arrow; scale bar, 2 μm.

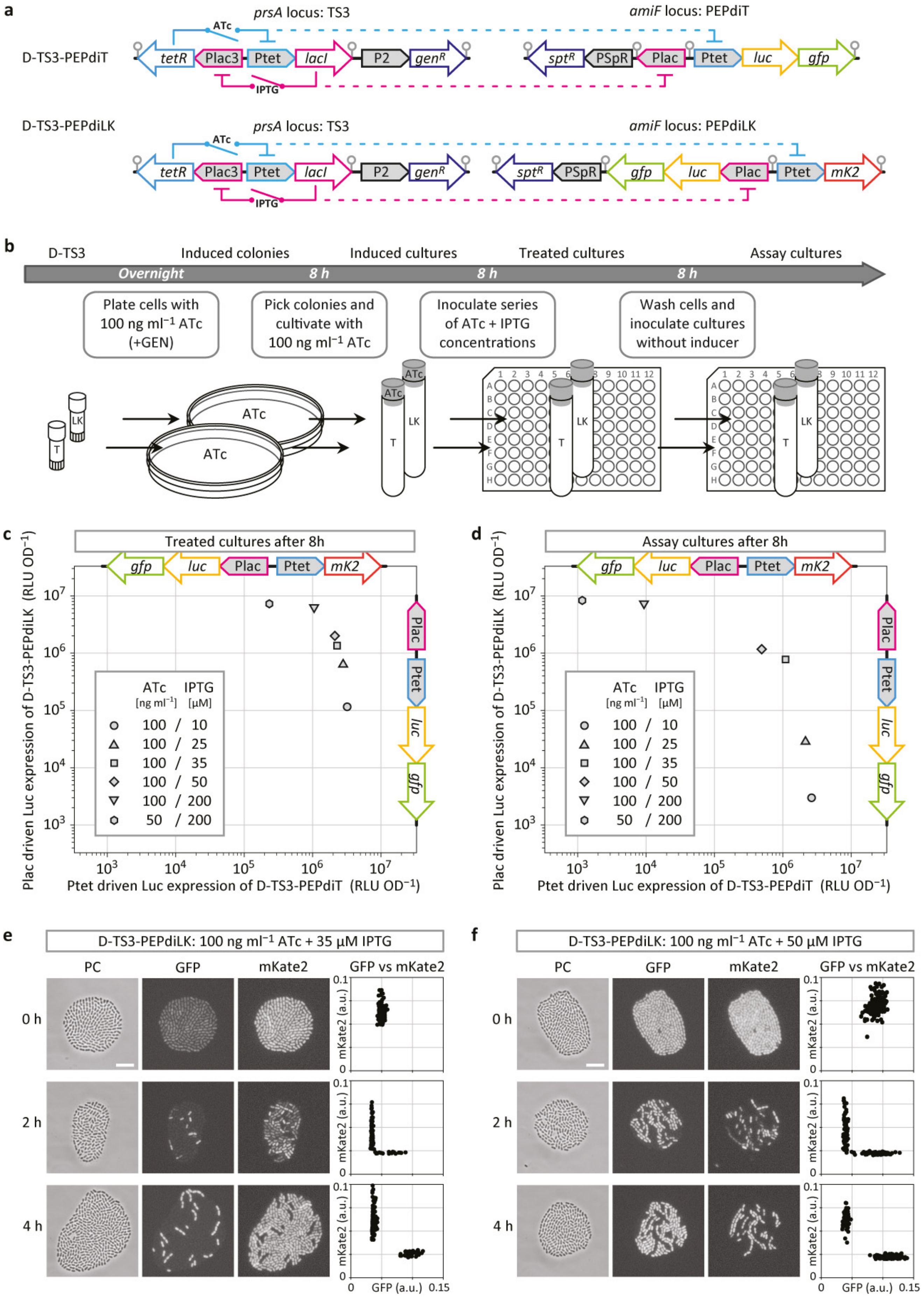
a
D-TS3-PEPdiT
prsA locus: TS3
amiF locus: PEPdiT
ATc
tetR
Plac3
Ptet
lacI
P2
genR
IPTG
sptR
PSpR
Plac
Ptet
luc
gfp
D-TS3-PEPdiLK
prsA locus: TS3
amiF locus: PEPdiLK
mK2
b
D-TS3
Induced colonies
Induced cultures
Treated cultures
Assay cultures
Overnight
8 h
8 h
8 h
Plate cells with 100 ng ml−1 ATc (+GEN)
Pick colonies and cultivate with 100 ng ml−1 ATc
Inoculate series of ATc + IPTG concentrations
Wash cells and inoculate cultures without inducer
c
Treated cultures after 8h
d
Assay cultures after 8h
ATc [ng ml−1] IPTG [µM]
100 / 10
100 / 25
100 / 35
100 / 50
100 / 200
50 / 200
Plac driven Luc expression of D-TS3-PEPdiLK (RLU OD−1)
Ptet driven Luc expression of D-TS3-PEPdiT (RLU OD−1)
e
D-TS3-PEPdiLK: 100 ng ml−1 ATc + 35 µM IPTG
f
D-TS3-PEPdiLK: 100 ng ml−1 ATc + 50 µM IPTG
PC
GFP
mKate2
GFP vs mKate2
0 h
2 h
4 h
mKate2 (a.u.)
GFP (a.u.)
0
0.1
0.15

**Supplementary Figure 3 | Identification of threshold induction conditions.** (**a**) Schematic representation of strain D-TS3-PEPdiT, harboring toggle switch 3 (TS3) at the *prsA* locus and reporter genes at the *amiF* locus; grey circles indicate transcription terminators. (**b**) Work flow of Ptet and Plac induction of TS3 strains and the subsequent settlement at equilibrium states; LK, strain D-TS3-PEPdiLK; T, strain D-TS3-PEPdiT; parallel assays were employed for 'treated cultures' and 'assay cultures', in microtiter plates for luminescence detection and in 5 ml tubes for sampling. (**c**) ATc- and IPTG-treatment (see inset) of the strains shown in **a**, measured by luminescence after 8 h of cultivation, with D-TS3-PEPdiLK (upper scheme) readings in the *y*-coordinate and D-TS3-PEPdiT (scheme to the right) readings in the *x*-coordinate. (**d**) Luminescence of previously treated cells (see inset), measured after 8 h of cultivation without inducer, with readings of D-TS3-PEPdiLK (upper scheme) in the *y*-coordinate and readings of D-TS3-PEPdiT (scheme to the right) in the *x*-coordinate. (**e**, **f**) Phase contrast and fluorescence microscopy of D-TS3-PEPdiLK cells expressing GFP from Plac and mKate2 from Ptet, previously induced with 100 ng $ml^{-1}$ ATc + 35 µM IPTG (**e**) and 100 ng $ml^{-1}$ ATc + 50 µM IPTG (**f**), after 0, 2 and 4 h of growth in liquid culture without inducer; scale bar, 8 µm; on the right, cell area-normalized GFP versus mKate2 fluorescence (a.u., arbitrary units; see Methods) of individual cells is shown.

**Supplemental Movie S1 legend**

Time-lapse experiment of D-TS3-PEPdiLK cells previously treated with 100 ng ml-1 ATc + 50 µM IPTG growing on a semi-solid surface without inducer; scale bar, 5 µm. Images were taken every 20 min. See Materials and Methods for more details.
